# On the early-life origins of vulnerability to opioid addiction

**DOI:** 10.1101/716522

**Authors:** Sophia C Levis, Brandon S Bentzley, Jenny Molet, Jessica L Bolton, Christina R Perrone, Tallie Z Baram, Stephen V Mahler

## Abstract

The origins and neural bases of the current opioid addiction epidemic are unclear. Genetics plays a major role in addiction vulnerability, but cannot account for the recent exponential rise in opioid abuse, so environmental factors must contribute. Individuals with history of early-life adversity (ELA) are disproportionately prone to opioid addiction, yet whether ELA interacts with factors such as increased access to opioids to directly influence brain development and function and cause opioid addiction vulnerability is unknown. We simulated ELA in female rats and this led to a striking opioid addiction-like phenotype. This was characterized by resistance to extinction, increased relapse-like behavior, and, as in addicted humans, major increases in opioid economic demand. By contrast, seeking of a less salient natural reward was unaffected by ELA, whereas demand for highly palatable treats was augmented. These discoveries provide novel insights into the origins and nature of reward circuit malfunction that may set the stage for addiction.

## Introduction

The devastating epidemic of opioid addiction in the U.S is enormously costly, in both human life and economically^1^. Genetics plays a major role in addiction vulnerability, but cannot account for this recent rise in opioid abuse, so environmental factors (e.g. increased access to prescription opioids) must contribute^2-4^. Notably, substance use disorders are most prevalent among those who experienced early-life poverty and trauma, and given their over-representation in this population, women are especially vulnerable^5-9^. However, it is yet unclear whether early-life adversity actually *causes* opioid use disorder vulnerability through alterations of the maturation and function of pleasure/reward circuits.

In humans, separating influences of ELA from the roles of genetics and other contributing factors is complex^10^. Therefore, we simulated resource scarcity in rats during a short, defined postnatal sensitive period by limiting bedding and nesting materials in the cage (LBN)^11, 12^. This leads to aberrant maturation of brain circuits underlying reward and stress^12^, which are crucially implicated in addiction to a range of abused drugs.

Here, we tested whether ELA promotes the use, seeking, and addiction to opioid drugs, and compared these behaviors to the seeking of natural rewards including highly palatable food rewards, all indicative of specific alterations in reward circuit function. We report here on a phenotype of several opioid-addiction-like behaviors in ELA experiencing female rats. This was characterized by resistance to extinction, increased relapse-like behavior, and, as in addicted humans, major increases in economic demand. By contrast, seeking of less salient natural rewards was unaffected by ELA, whereas demand for highly palatable food was also augmented.

## Materials and Methods

### Animals

Primiparous, timed-pregnant Sprague-Dawley rats were obtained from Envigo (Livermore, CA) on E15, and maintained in an uncrowded, quiet animal facility room on a 12 h light/dark cycle. Parturition was checked daily, and the day of birth was considered postnatal day (PD) 0. On PD2, litters were mixed and both male and female pups were assigned to ELA and control (CTL) groups. Rats were weaned at PD21, and females were then housed in groups of 2-4, with a 12 h reverse light cycle. Food and water were available *ad libitum* throughout all experiments. All procedures were approved by the University of California Irvine Institutional Animal Care and Use Committee and conducted in accordance with the National Institutes of Health guide for the care and use of laboratory animals.

### The Limited Bedding and Nesting (LBN) model of early-life adversity

On PD2, pups from at least two litters were gathered, and pups were assigned at random to each dam in equal numbers of male and female to prevent potential confounding effect of genetic variables or litter size. Dams and pups assigned to the LBN group were transferred to cages fitted with a plastic-coated mesh platform sitting ∼2.5□cm above the cage floor. Bedding sparsely covered the cage floor under the platform, and one-half of a 24.2□cm□ ×□23.5□cm paper towel was provided for nesting material. Control group (CTL) dams and pups were placed in cages containing a standard amount of bedding (∼0.33 cubic feet of shredded corn cob) without a platform, and one full paper towel. CTL and LBN cages remained undisturbed during PD2-9, during which maternal behaviors were monitored as previously described^13-15^. On PD10, animals were all transferred to CTL condition cages.

### Intravenous catheter surgery

At approximately PD70, rats were deeply anesthetized with isoflurane (2-2.5%) and chronic indwelling catheters were inserted into the right jugular vein, exiting two cm caudal to the scapulae. Meloxicam (1mg/kg, i.p.) for postsurgical analgesia, and prophylactic antibiotic cefazolin (0.2ml, i.v.; 10mg/0.1ml) were given perioperatively. After 5 days of recovery, catheters were flushed daily following each opioid self-administration session with cefazolin (10mg/0.1ml) and heparin lock solution (10 U/0.1ml) to maintain catheter patency.

### Drugs

Heroin (diamorphine) HCl was provided by the National Institute on Drug Abuse (NIDA) Drug Supply Program (Research Triangle Park, NC, USA) and Cayman Chemical Company (Ann Arbor, MI, USA), and remifentanil HCl was provided by the NIDA Drug Supply Program. All drugs were dissolved in sterile 0.9% saline for experimental use. Animals self-administered intravenous heroin at a concentration of 0.1mg/kg/infusion during acquisition (days 1-3), and 0.05mg/kg/infusion during maintenance of heroin self-administration. For the remifentanil behavioral economic task, remifentanil was self-administered at decreasing concentrations over successive 10-minute bins, as follows: 1.92, 1.08, 0.608, 0.342, 0.192, 0.108, 0.0609, 0.0341, 0.0193, 0.0108, 0.00611 µg remifentanil/infusion.

### Behavioral / Functional Tests

M&M consumption, heroin self-administration, extinction, reinstatement, and remifentanil demand tests were all performed in the same female rats (n=16). Sucrose preference tests were performed in a separate cohort of females (n=24). To further examine the effects of ELA on economic demand, female rats from a third cohort underwent, sequentially, both food (chow and palatable), and then remifentanil demand tasks (n=17). Sample sizes were chosen based on observed effect sizes in our prior reports^11-14^.

### Sucrose Preference

Adult ELA and CTL females underwent a two-bottle home cage sucrose preference test^14^. The test consisted of two phases: one week of habituation to two bottles containing 50 mL tap water, followed by two weeks of sucrose consumption. In the second phase, one of the bottles was switched to contain 50 mL of 1.5% sucrose. The left/right position of the bottles was counterbalanced to obviate side bias. Fluid consumption was measured each morning, and drinking solution was refreshed to 50 mL daily.

### M&M Consumption

Rats were given a three-day test for free intake of M&M brand chocolates (Mars, McLean, VA) prior to catherization and opiate self-administration training. On three consecutive days, animals were habituated to tub cages for one hour, and then given ad libitum access to chocolate M&M’s for one hour. M&M consumption (g) was totaled over all three days for analysis.

### Acquisition and Maintenance of Heroin Self-Administration

Self-administration training and testing took place in Med Associates operant chambers within sound-attenuating boxes, each equipped with two retractable levers with white lights above them, a white house light, and a tone generator. Intravenous heroin was administered via a pump located outside the sound-attenuating box. Rats received daily 2-hr self-administration sessions, when pressing on the “active” lever yielded a heroin infusion of 0.1 mg/kg (acquisition days 1-3) or 0.05 mg/kg (maintenance days 4-17), achieved by varying infusion volume based on animals’ weight. Heroin infusions were accompanied by concurrent 2.9-kHz tone and lever light illumination for 3.6s (fixed ratio 1 schedule: FR1). A 20-sec timeout period (signaled by turning off the house light) followed each infusion/cue presentation, during which additional lever presses were recorded but had no consequence. Pressing on the second “inactive” lever was recorded but had no consequence. Doses were chosen that have previously been shown to sustain self-administration^16^.

### Heroin Extinction and Reinstatement

Following heroin self-administration, rats received extinction training for a minimum of seven days, or until an extinction criterion of two consecutive days of less than 20 active lever presses was met. During extinction training, presses on both levers were recorded but lever pressing had no consequence. Upon meeting the extinction criterion, rats underwent a cue-induced reinstatement test, during which one presentation of the previously drug-paired light and tone cue was delivered 10s after the start of the session, then presses on the active lever yielded additional cue presentations as in self-administration training, but no heroin. Next, rats underwent a minimum of 2 additional extinction training days until extinction criterion was re-attained, after which they underwent a heroin-primed reinstatement test. The rats received an experimenter-administered injection of heroin (0.25mg/kg, s.c.^16^), or saline vehicle, immediately prior to being placed into the operant boxes for a 2-hr session. Lever presses were recorded but did not yield additional heroin or cues (extinction conditions). All animals received both heroin and vehicle-primed reinstatement tests on separate days in counterbalanced order. Again, 2+ additional extinction training days occurred between primed reinstatement tests to re-establish extinction criterion. Following reinstatement tests, catheter patency was confirmed by administering intravenous brevital (0.1-0.2ml, 5mg/ml). Rats with catheter failure were re-catheterized in the left jugular vein, and allowed to recover prior to starting behavioral economic thresholding procedures.

### Behavioral economic thresholding procedure

Rats were trained on a previously-described within-session economic thresholding procedure^17-20^. In order to avoid undesired effects of drug satiety on responding due to the long half-life of heroin, the short-acting opioid remifentanil, a fentanyl analog, was used for this task. On each training day, presses on the active lever delivered remifentanil and cue presentations on an FR1 schedule. The duration of each cue/infusion was decreased every 10 min across the 110-min session, requiring rats to exert increasing effort to maintain a desired blood level of drug^19, 20^. Drug intake was determined at the various response requirements (i.e. costs) and consumption data was then modeled with an exponential demand equation 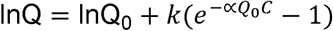 as previously described^17-19^, yielding variables corresponding to hedonic set point (Q_0_, reflecting extrapolated value of remifentanil at price 0) and motivation (α, reflecting demand elasticity). Rats were run daily on the threshold procedure for 14 days, and behavior derived from the average of the last 5 days was analyzed as described above.

### Palatable and nutritive food thresholding procedure

Rats were trained on a modified economic thresholding procedure^21^ to assess demand for highly palatable food pellets containing sucrose (52%), fat (6.3%), and protein (20.2%), and normal chow pellets (fat: 5.6%; fiber: 4.7%; protein: 18.7%) as a less salient food reinforcer. Rats initially underwent daily 1hr training sessions with an increasing fixed-ratio (FR) schedule between days (FR1, 3, 10, 32, 100). Rats moved to a higher FR only after having reached responding criteria (criteria: FR1 >60 pellets, FR3 >20 pellets, FR10 >6 pellets, FR32 >2 pellets, FR100- no pellet requirement). All animals were trained up to FR100 prior to moving on to thresholding task. Rats were then trained on a reverse thresholding procedure in which the number of lever presses yielding a food pellet and light/tone cue decreased incrementally every 25 minutes (5 minutes access to lever-pressing followed by 20-minute timeout) throughout the 105-minute session (FR100, 32, 10, 3, 1). Rats were trained on the thresholding procedure for 10 days each for palatable food, then chow, and performance on the last 5 days of training for each food reinforcer was fit to a demand curve and analyzed using the exponential demand equation described above.

### Analysis and Statistics

Independent samples t-tests were used to determine effects of ELA on sucrose preference, M&M consumption, heroin consumption, remifentanil low-effort dose preference, days until extinction criterion, reinstatement, and demand (Q_0_, α) for remifentanil, and palatable/non-palatable foods. Extinction persistence was further analyzed for “survival’ of seeking behavior (i.e. percentage of rats in each group failing to meet extinction criterion on each training day), and plots were compared using the log-rank test^22^. For analyses containing within-subjects variables (ex. saline vs. heroin reinstatement), mixed model ANOVAs were used, with group (ELA/CTL) as a between-subjects factor. Following significant interactions in ANOVA, Bonferroni post-hoc tests were used to characterize the nature of effects. Randomization was not used in the design of these studies, and experimenters were not blind to experimental group during testing. Compared groups did not statistically differ from one another in variance, accommodating assumptions of the parametric tests employed.

## Results

### Heroin Self-Administration, Extinction, and Reinstatement

We found strikingly augmented addiction-like behaviors for two distinct classes of opioid drugs, heroin and remifentanil, in adult female rats raised under ELA conditions. Total heroin self-administration did not vary between control and ELA rats (**Fig. 1A;** CTL mean(SEM) = 13.01(0.9687), ELA m = 11.37(0.4302); t_12_=1.550; P=0.1471), nor did responding for remifentanil at low effort during the demand task (mean µg/kg taken in first 10min of the demand sessions: CTL = 41.79, ELA = 43.06; t_29_ = 0.5844; P = 0.5634; **Fig. 1B).** However, once heroin was withdrawn after self-administration training, the ELA group resisted extinction, a robust indicator of compulsive drug seeking^23, 24^ (**Fig. 1C;** mean (SEM) days until criterion; CTL m = 5.714(0.4738), ELA m = 8.429(0.9724); t_12_=2.509; P=0.0274). Although initial responding on extinction training day 1 was unaltered (no group X 15min time bin interaction for active lever pressing: F_(7,84)_ = 1.482; P=0.1846), the probability that ELA rats will achieve extinction criterion each day was lower than in controls (**Fig. 1D**; Kaplan-Meier probability of survival; CTL median survival = 6 days; ELA median survival = 8 days; Log-rank curve comparison Chi square (df)= 4.491(1), P = 0.0341). Moreover, post-extinction reinstatement of heroin seeking elicited by either heroin cues or a priming dose of heroin itself was greater in ELA rats compared with CTLs; **Fig. 1E,F**). Notably these reinstatement measures are considered risk factors for craving and relapse to drug use in humans attempting to quit.^25, 26^ Significant cue-induced and heroin-primed reinstatement was apparent in all rats (main effect of cue/no-cue: F_(1,12)_ = 89.20; P < 0.0001; main effect of heroin/saline prime: F_(1,12)_ = 27.00; P = 0.0002). However, both types of reinstatement were augmented in ELA animals (Bonferroni’s post-hocs; cue: t_24_ = 4.676; P=0.0002; prime: t_24_ = 3.210; P=0.0075).

**Figure 1:**
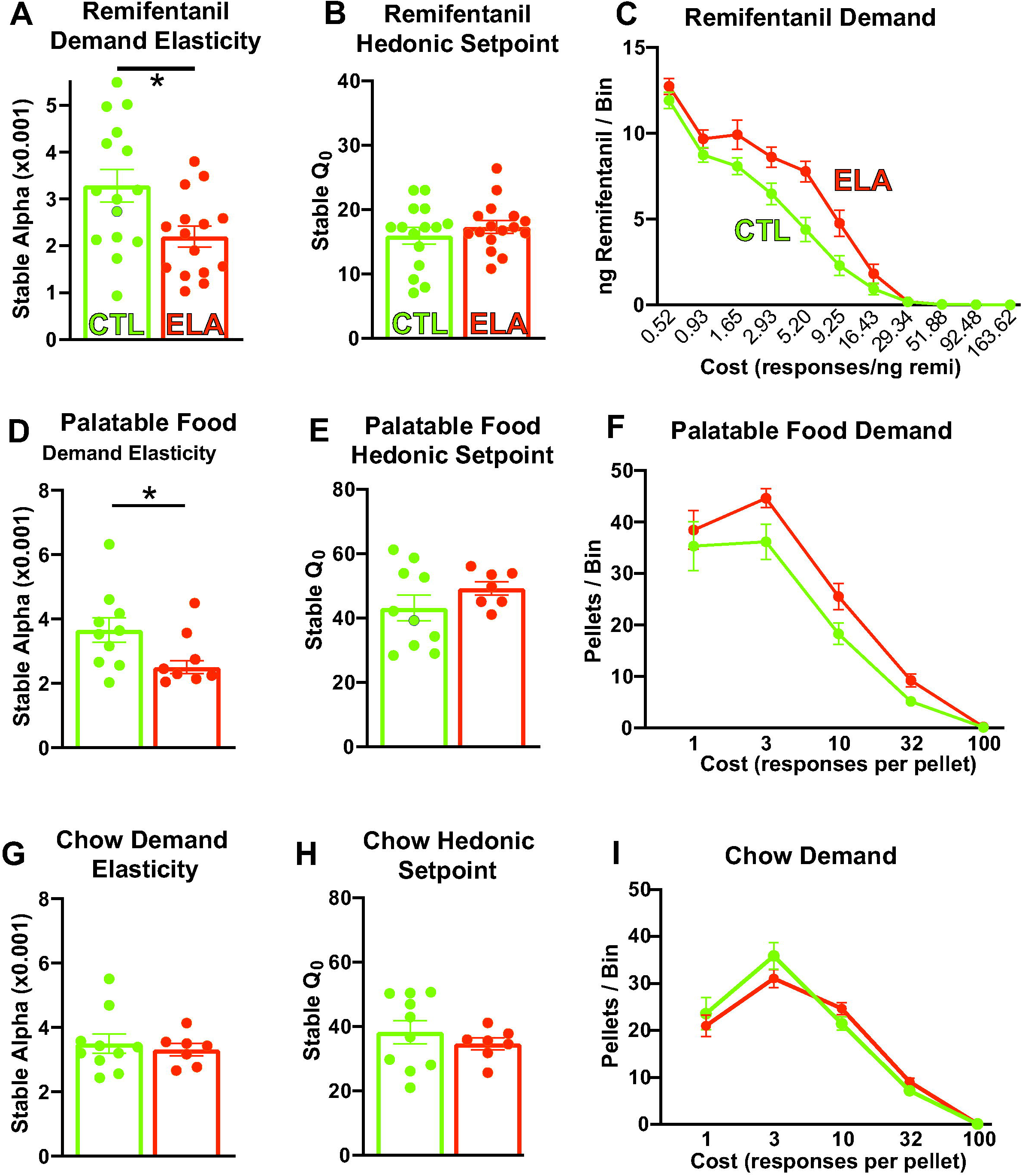
Early-Life Adversity Potentiates Heroin Seeking Behaviors: (A) Total heroin self-administration did not differ between control and ELA rats (t_12_=1.550; P=0.1471), nor did (B) remifentanil self-administration at low effort (t_29_ = 0.5844; P = 0.5634). (C) Once response-contingent heroin was withdrawn, the ELA group resisted extinction more than controls (t_12_=2.509; P=0.0274). (D) Accordingly, the probability that ELA rats achieve extinction criterion (<20 active lever presses) each day was lower than in controls (Kaplan-Meier probability of survival; Log-rank curve comparison Chi square(df) = 4.491(1), P = 0.0341). (E,F) ELA augmented reinstatement of heroin-seeking by heroin-associated cues (main effect of cue/no cue: F_(1,12)_ = 89.20; P<0.0001; main effect of rearing condition: F_(1, 12)_ = 9.982; P = 0.0082; cue x ELA interaction: F_(1,12)_ = 12.49; P = 0.0041. Bonferroni’s post-hoc tests CTL vs ELA: no cue: t_24_ = 0.2543; P > 0.9999; cue: t_24_ = 4.676; P = 0.0002), and by a heroin priming injection (main effect of heroin/saline prime: F_(1,12)_ = 27.00; P = 0.0002; main effect of rearing condition: F_(1, 12)_ = 4.891; P = 0.0471; prime x ELA interaction: F_(1,12)_ = 5.447; P = 0.0378. Bonferroni’s post-hoc tests CTL vs ELA: saline prime: t_24_ = 0.01554; P > 0.9999; heroin prime: t_24_ = 3.210; P = 0.0075). Circles within each bar represent individual animals in CTL (green) and ELA (orange) groups. *p<0.05; **p<0.01; ***p<0.001.

### Demand for Remifentanil

In humans, drug addiction is characterized by high demand for drugs that is insensitive to escalating financial, social, and health costs. This phenomenon, termed ‘demand inelasticity,’ is characteristic of economic demand for a wide range of essential commodities, and is a hallmark of compulsive drug seeking in people with substance use disorders^27, 28^. Therefore, we tested demand elasticity for an opioid drug using an established and sensitive assay of microeconomic demand^17, 18, 29^. Because this within-session protocol requires short-acting drugs which enable robust lever-pressing throughout a ∼2hr session^19^, we examined demand for the short-acting fentanyl analog remifentanil in control and ELA rats. This approach additionally broadened the range of drugs tested to a distinct class of abused opioids.

Remarkably, ELA-reared rats exhibited features similar to human opioid addicts^30^ when given the opportunity to self-administer remifentanil. Specifically, they were willing to pay a significantly higher price to obtain the drug compared to controls—their demand became inelastic and relatively insensitive to cost (decreased α) (**Fig. 2A**; CTL mean(SEM) = 3.286(0.3502), ELA mean(SEM) = 2.193(0.2237); t_28_=2.630; P=0.0137). The selectivity of this addictive-like behavior was striking: no changes were observed in the preferred intake of remifentanil by the ELA group when the drug was essentially “free”, measured as the hedonic set-point, or Q_0_ (**Fig. 2B**; CTL mean(SEM) = 15.97(1.307), ELA mean(SEM) 17.32(1.011); t_28_=0.8159; P=0.4215). Accordingly, the total amount of remifentanil taken under low effort conditions in these tests did not differ between the CTL and ELA groups (**Fig. 1B**). These findings align with our finding that low effort heroin intake did not differ between groups (**Fig. 1A**). Average drug intake in each effort bin is shown in **Fig. 2C**.

**Figure 2:**
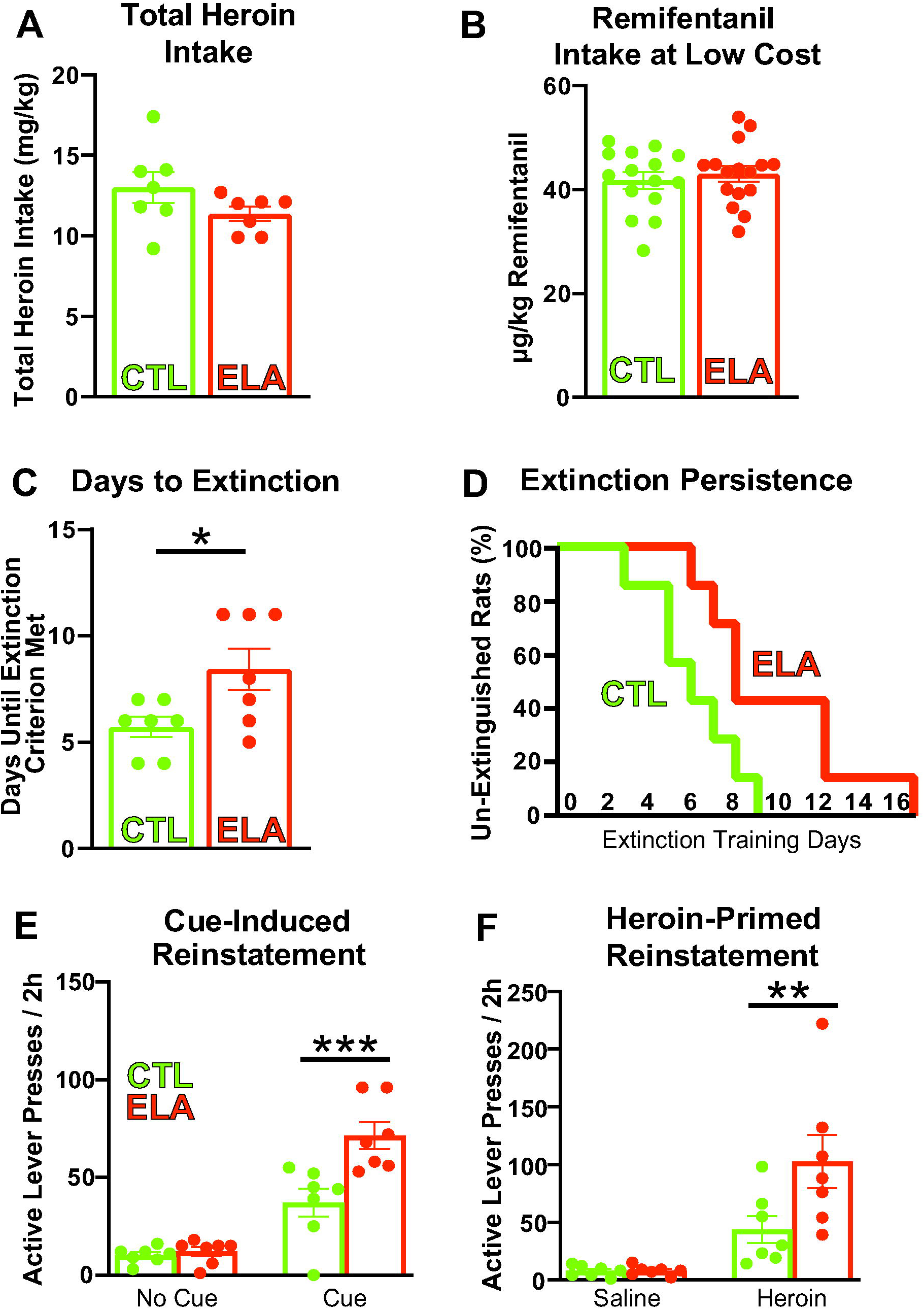
Early-Life Adversity Increases Demand for Opioids and Highly Palatable Food: (A) ELA-reared rats had a decreased sensitivity to cost, or an ‘inelastic’ demand, for remifentanil compared to controls (t_28_=2.630; P=0.0137). (B) However, no changes were observed in the intake of remifentanil by the ELA group when the drug was essentially “free”, measured as the hedonic set-point, or Q_0_ (t_28_=0.8159; P=0.4215). (C) Average (±SEM) drug intake at each effort step is depicted. (D) ELA rats were willing to pay a significantly higher price (effort) for a highly palatable food, indicating decreased demand elasticity, or α (t_15_=2.341, P=0.0334), whereas (E) consumption at low effort (Q_0_) did not differ between ELA and CTL rats (t_15_=1.199, P=0.2490). (F) Average palatable food intake at each effort step is depicted. (G) In contrast to palatable food, no difference was seen in demand elasticity (α) between ELA and CTL rats for chow pellets (t_15_=0.4619, P=0.6508), and (H) Consumption at low effort (Q_0_) also did not differ between ELA and CTLs (t_15_=0.7922, P=0.4406). (I) Average chow pellet intake at each effort step is depicted. Circles within each bar represent individual animals in CTL (green) and ELA (orange) groups. *p<0.05.

### Palatable and Nutritive Food Reward

In a typical sucrose preference assay of natural reward sensitivity, or “anhedonia”^31-33^, ELA females were indistinguishable from those reared in control lab conditions, unlike prior findings in males^12^. Percent of sucrose consumption (vs. water) on the two-bottle task did not differ based on early-life rearing conditions (**Supplemental Fig. 1A**; CTL m = 75.19%(1.729), ELA m = 76.42%(1.221); t_22_=0.5831; P=0.5657). In contrast, ELA females ate more highly-palatable M&M chocolates than their CTL counterparts in a separate assay of highly palatable reward intake^17^ (**Supplemental Fig. 1B;** CTL m = 3.701g(0.2763), ELA m = 4.852g(0.2121); t_14_=3.365; P=0.0046).

Next, we examined economic demand for highly palatable food morsels and typical nutritive chow in ELA and CTL rats. ELA rats were willing to pay a significantly higher price (in effort) for the palatable food pellets, indicating decreased demand elasticity (**Fig. 2D**, CTL α m = 3.660(0.3865), ELA α m = 2.503(0.1978); t_15_=2.341, P=0.0334). In contrast, when the rats worked for chow pellets, no difference was seen in demand elasticity between ELA and CTL rats (**Fig 2G**, CTL α m = 3.498(0.2970), ELA m = 3.317(0.1909); t_15_=0.4619, P=0.6508). Measures of baseline intake (lever presses resulting in food at low effort Q_0_) did not differ between ELA and CTLs for either palatable food pellets or chow (**Fig. 2E,H;** palatable food: CTL m = 43.10(3.974), ELA m = 49.22(2.113), t_15_=1.199, P=0.2490; chow: CTL m = 38.25(3.599), ELA m = 34.60(1.845), t_15_=0.7922, P=0.4406). Average food pellet consumption in each effort bin is shown in **Figs. 2F,I**.

## Discussion

Together, our findings indicate that in a preclinical experimental system where genetics and prior existing conditions can be controlled, poverty-like rearing during sensitive developmental periods^34, 35^ can lead to vulnerability to the addictive effects of opioid drugs. Indeed, the findings suggest that critical developmental periods—such as those which are well-defined for sensory circuits^34, 35^—apply also to developing reward circuitries, and potentially to other cognitive and emotional brain systems as well^11, 12, 14, 36^.

The present results reveal that the consequences of aberrant reward circuit development caused by ELA may differ between males and females. For example, ELA in females here led to exaggerated pursuit of heroin and remifentanil, coupled with increased motivation for highly-palatable food (sweet banana pellets or chocolate), without impacting intake of less salient food rewards (chow pellets or a dilute sucrose solution). In contrast, we previously showed in males that ELA induces persistent anhedonia for food (sucrose preference; M&M intake) and social play^12^. Males also exhibited a lower hedonic set-point to an addictive drug in the demand elasticity test, though the drug was cocaine rather than an opioid^17^. We note that the remarkable ability of ELA to increase motivation and demand for highly hedonic, salient rewards specifically in females may help explain the prevalence of ELA in heroin-addicted women^6^.

Notably, our findings suggest that the influence of ELA on the normal maturation of reward circuits may differ in males and females, potentially affecting specific circuit nodes or molecularly defined pathways. Our findings highlight the importance of attention to sex differences in future mechanistic studies of ELA-related opioid addiction, and in tests of prevention or intervention strategies for this problem.

The specific aspects of early-life adversity that cause these effects, and the mechanisms by which ELA generates addiction-like behavior, remain to be explored. Our prior work demonstrated that ELA (using our naturalistic LBN model) is characterized by chaotic, unpredictable maternal signals to the developing pups^31^, and converging evidence from both humans and rodents shows that such unpredictable, fragmented early-life environments profoundly impact brain function later in life^32^—i.e. into adolescence^14, 37, 38^ and persisting into adulthood^12, 17, 39^. A potential mechanism for these long-term negative outcomes, including opioid addiction risk, is aberrant maturation of specific reward- and stress-related brain circuits, a finding that also emerges in studies using translatable methods such as structural magnetic resonance imaging^11^.

Whereas numerous aspects of addiction vulnerability we have discovered here require further investigation (including sensitivity to opiate drug dosages^40, 41^), the present work sets the groundwork for mechanistic studies of the responsible alterations in reward circuit functions. Importantly, it creates for the first time a direct causal link between ELA and opioid addiction vulnerability, and establishes a new framework for testing mechanisms across species, ultimately enabling the development of predictive biomarkers and prevention.

## Acknowledgements

We thank the National Institute on Drug Abuse (NIDA) Drug Supply Program (Research Triangle Park, NC, USA), for providing heroin and remifentanil. The research was supported in part by NIH grants MH096889; NS28912; DA035251; GM008620; MH119049, and the Hewitt Foundation for Biomedical Research

## Data availability

The data forming the basis of these studies do not include genomic or imaging data sets storable in public repositories. Data will be readily made available when requested from the authors.

