## Supplementary figures and images for "On the early-life origins of vulnerability to opioid addiction"

### Supplemental Figure 1

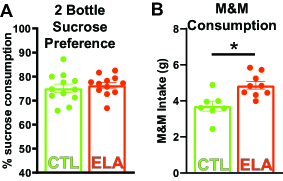
